# High-density super-resolution microscopy with an incoherent light source and a conventional epifluorescence microscope setup

**DOI:** 10.1101/132571

**Authors:** Kirti Prakash

## Abstract

We report that single-molecule superresolution microscopy can be achieved with a conventional epifluorescence microscope setup and a Mercury arc lamp. The configuration termed as Omnipresent Localisation Microscope (OLM), is an extension of Single Molecule Localisation Microscopy (SMLM) techniques and allows single molecules to be switched on and off (’blinking’), detected and localised. The use of a short burst of deep blue excitation can be further used to reactivate the blinking, once the blinking process has slowed or stopped. A resolution of 90 nm is achieved on test specimens (mouse and amphibian meiotic chromosomes). Finally, for the first time, we demonstrate that STED and OLM can be performed on the same biological sample using a simple imaging buffer. It is hoped that such a correlative imaging will provide a basis for a further enhanced resolution.

**Scope of the findings:** Despite ten years of development, superresolution microscopy is still limited to relatively few microscopy and optics groups. This is mainly due to the significant cost of the superresolution microscopes which require high-quality lasers, high NA objective lens, a very sensitive camera, a highly precise microscope stage, and a complex post-acquisition data reconstruction and analysis. We present results that demonstrate the possibility to obtain nanoscale resolution images using a conventional microscope and an incoherent light source. We show an easyto-follow protocol that every biologist can implement in the laboratory. We hope that this finding will help any scientist to generate high-density super-resolution images even with limited budget. Lastly, the new photophysical observations reported here will pave the way for more in-depth investigations on excitation, photobleaching and photoactivation of a fluorophore.

---

Resolution of images generated by light microscopy is limited by diffraction due to the wave nature of light (Born and Wolf, 1980; Betzig and Trautman, 1992; Cremer and Masters, 2013). As a consequence, in the image space, a point-like object is registered as a blurred Airy-disc. The function describing this blurring is referred to as the Point Spread Function (PSF), and can be described in terms of the numerical aperture (NA) of the objective lens and the wavelength (*λ*) of the light used. When the most optimal combination of objective lenses and light in the visible region is used, a resolution of about 200 nm can be achieved, which is often referred to the resolution limit of light microscopes (Abbe, 1873; Rayleigh, 1896).

The resolution limit of light microscopes has been recently overcome by a number of superresolution microscopy techniques (SMTs), such as single molecule localization microscopy (SMLM) (Lidke et al., 2005; Betzig et al., 2006; Hess et al., 2006; Rust et al., 2006), stimulated emission depletion (STED) (Hell and Wichmann, 1994; Willig et al., 2006), structured illumination microscopy (SIM) (Heintzmann and Cremer, 1999; Gustafsson, 2000) and various related techniques (Heilemann et al., 2008; Schoen et al., 2011; Szczurek et al., 2014; Dertinger et al., 2009; Gustafsson et al., 2016).

Currently, all the superresolution microscopy techniques (SMTs) make use of coherent light sources such as lasers. For most SMTs, lasers are a necessity, for instance, STED. For single molecule based superresolution techniques a minimum power (approximately 0.1kW/cm2) is required to switch the fluorophores between dark and bright state, often termed as ‘blinking.’ We wondered if one can make use of an incoherent light source such as a lamp instead of lasers to induce the on/off switching of single molecules. Previously, it has been shown that incoherent light sources such as Mercury arc lamps and LEDs can be used for detection of single molecules (Chiu and Quake, 1999; Gerhardt et al., 2011). In this report, we demonstrate that using a Mercury arc lamp and appropriate combination of dyes and filters, one can extract enough photons to detect, localise and reactivate a fluorophore with a nanometer precision.

We applied a simple protocol on a standard epifluorescence microscope setup to generate high-resolution images of synaptonemal complex and lampbrush chromosomes and compared them with other state-of the-art superresolution techniques. In this configuration, single fluorescent molecules undergo repeated cycles of fluorescent bursts, similarly to what can be observed with other SMLM technologies. The individual blinking events are detected with a CCD camera and localised with high precision. The positions of individual molecules are then used to reconstruct an image with high spatial resolution. The novel features are the following:

- On/off switching of single molecules by a Mercury arc lamp instead of coherent light sources such as lasers. Most super-resolution methods currently use lasers.
- High-resolution single-molecule images with an unmodified epifluorescence microscope, as found in all cell biology laboratories, instead of a specially constructed instrument.
- The use of a short burst of deep blue excitation to reactivate blinking, once blinking has slowed or stopped. Prolongated blinking helped to reconstruct super-resolved images of high signal density.
- Ability to perform STED and SMLM measurements on the same biological sample employing a simple imaging buffer (ProLong Diamond).

Regarding the different observations listed above, and with the growing interest of the community for validation of experiments at the nanoscale, we believe that the results presented in this paper potentially have a very broad application.

## Results

### Description of the OLM setup

Most of the cell biology laboratories today are equipped with an epifluorescence microscope setup (Figure 1A), for which the Mercury arc lamp is a common source of illumination. The Mercury arc lamp emits light with several peaks around 400 nm and 560 nm (Figure 1B). In fact, many dyes have their absorption spectra corresponding to the peaks of the Mercury arc lamp for optimal fluorescence. One such dye is Texas Red, which has its excitation peak around 594 nm. We used Alexa Fluor 594, a direct replacement of Texas Red, for its brighter signal and photostability. Alexa Fluor 594 can also be readily multiplexed with other fluorophores such as Alexa Fluor 405, Alexa Fluor 488, Alexa Fluor 647 for multicolour experiments.

**Figure 1.**
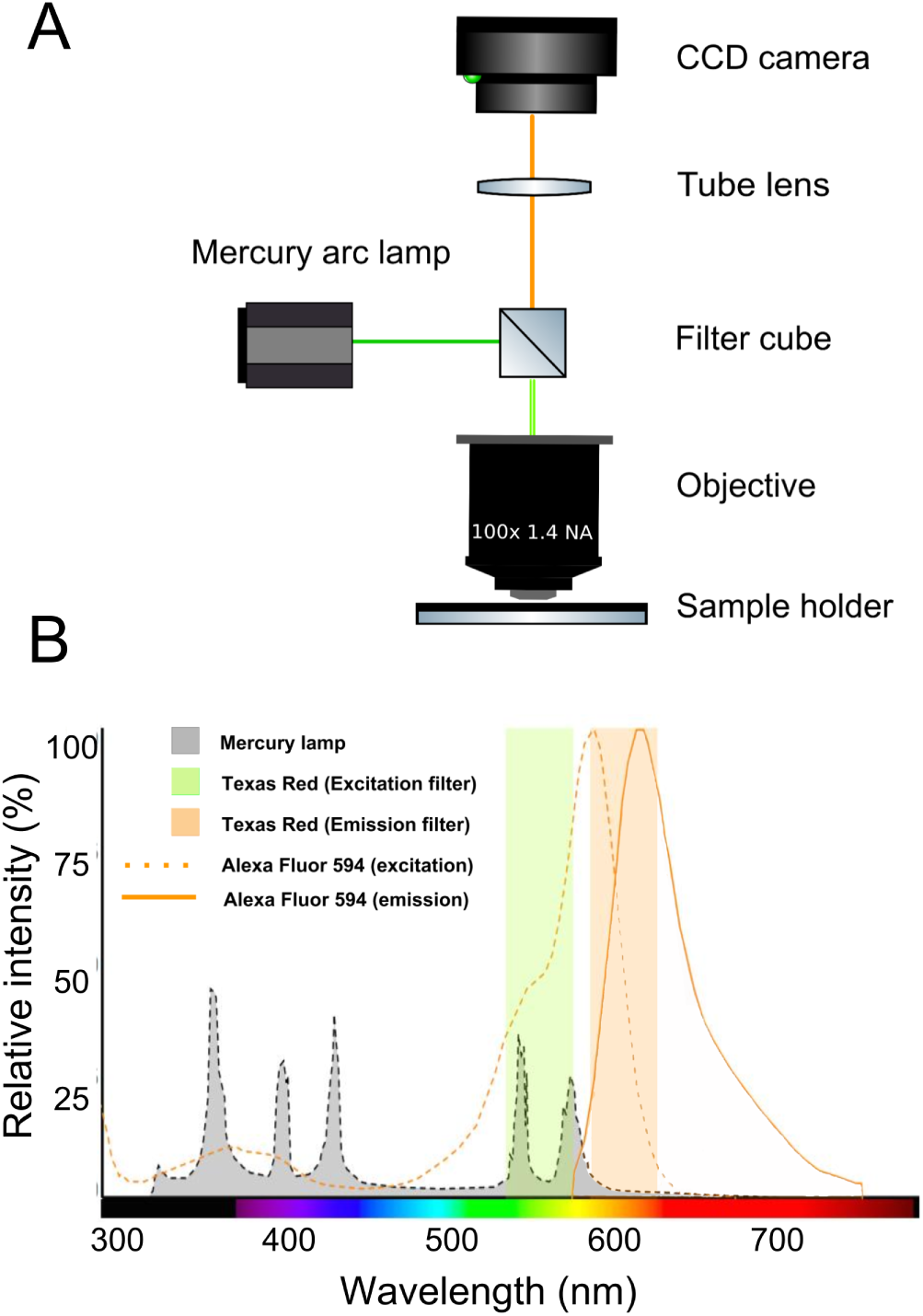
The OLM setup: (A) Schematic of the optical system of a standard epifluorescence microscope, as used in OLM. (B) The shaded grey area shows the emission spectrum of a Mercury arc lamp. We use the two peaks around 546 nm and 579 nm to bring the fluorophore (Alexa 594) to the temporary dark state. The fluorophore then begins to blink, emitting light with a maximum around 620 nm.

For measurements, we used Olympus BX61 (Figure S4), a standard widefield setup. The microscope was equipped with a Mercury arc lamp, 1.40 N.A., a 100x objective lens and a digital CCD camera. In the illumination module, we used Texas Red excitation filter (560/40 nm), which transmits all wavelengths between 546 and 579 nm peaks of the Mercury lamp (Figure 1B).

We start to record images once the individual molecules start to blink after the initial bleaching step (approximately 10-20 minutes, but the time depends on the antibody concentration). For Alexa Fluor 594, 0.2 kW/cm2 (Texas Red filter) of Mercury arc lamp power in the sample plane was sufficient to excite the activated molecules. After 1-2 min of imaging with Texas Red filter, the blinking diminishes and one needs to switch to the DAPI filter (365/30 nm, 0.05 kW/cm2 in the sample plane) to reactivate the molecules from the dark state to the bright state. The process of photobleaching and subsequent photoactivation for a sparse set of molecules needs to be repeated (approximately 1 to 2 hours) until sufficient signal density is reached for high-resolution image reconstruction.

To test the setup, we studied various kinds of samples that varied in thickness and spacing of molecular components: Synaptonemal Complex (SC) in mouse oocytes, Nuclear Pore Complexes (NPCs) and Lampbrush Chromosomes (LBCs) in amphibian oocytes. One of the main challenges of widefield illumination is that it lacks optical sectioning capabilities of confocal (Sheppard and Wilson, 1981) and multiphoton microscopy (Zipfel et al., 2003). Often TIRF objectives or special illumination strategies such as HILO or light sheet illumination are used to address the poor optical sectioning capability of widefield illumination (van’t Hoff et al., 2008; Keller et al., 2010). The flat NPC (approx. 100 nm in thickness) and SC (approx. 200 nm in thickness) samples helped us to circumvent the use of expensive TIRF objective lens, while imaging thick lampbrush chromosomes (approx. 1*µ*m), we removed the out-of-focus signals during the post-processing of the data based on the width of the PSF. Nonetheless, thick samples (LBCs) and samples with closely spaced molecular components (NPCs) remain a challenge for OLM.

On the microscopy side, use of the Mercury arc lamp helped to avoid the usual problems associated with lasers: alignment, need for multiple spectral lines, expense, danger to the eyes and premature bleaching of the fluorophores. Finally, use of ProLong Diamond as the imaging buffer further minimised the effort to prepare a complex cocktail of reducing/oxidising reagents to create a redox environment (Heilemann et al., 2008; Löschberger et al., 2012).

### On/off switching of single molecules induced by OLM

To test if individual molecules can be switched on/off with an incoherent light source, we labelled gp210, a protein known to occupy the nuclear pore complex periphery, with Alexa Fluor 594. At first, the dye fluoresces as usual. Over a matter of minutes the sample gradually bleaches entirely, but then individual molecules of fluorophore begin to ‘blink.’ That is, they fluoresce for a short time (seconds), then cease to fluoresce, and then fluoresce again. The blinking events are recorded with a relatively inexpensive CCD camera.

To quantify on/off switching of single molecules, we chose an area of 7X7 pixels (roughly 450X450 *nm*^2^) and plotted the intensity profile along the image stack (Figure 2A-C). We could observe sharp peaks, which are fluorescent bursts of single molecule integrated over 150 ms of camera exposure. The broad peaks in the profile occur when a fluorophore remains ‘on’ for an extended period. As we have no way to isolate such fluorophores optically, we merged such signals in all the consecutive frames during the final analysis (see Methods and Material section for more details).

**Figure 2.**
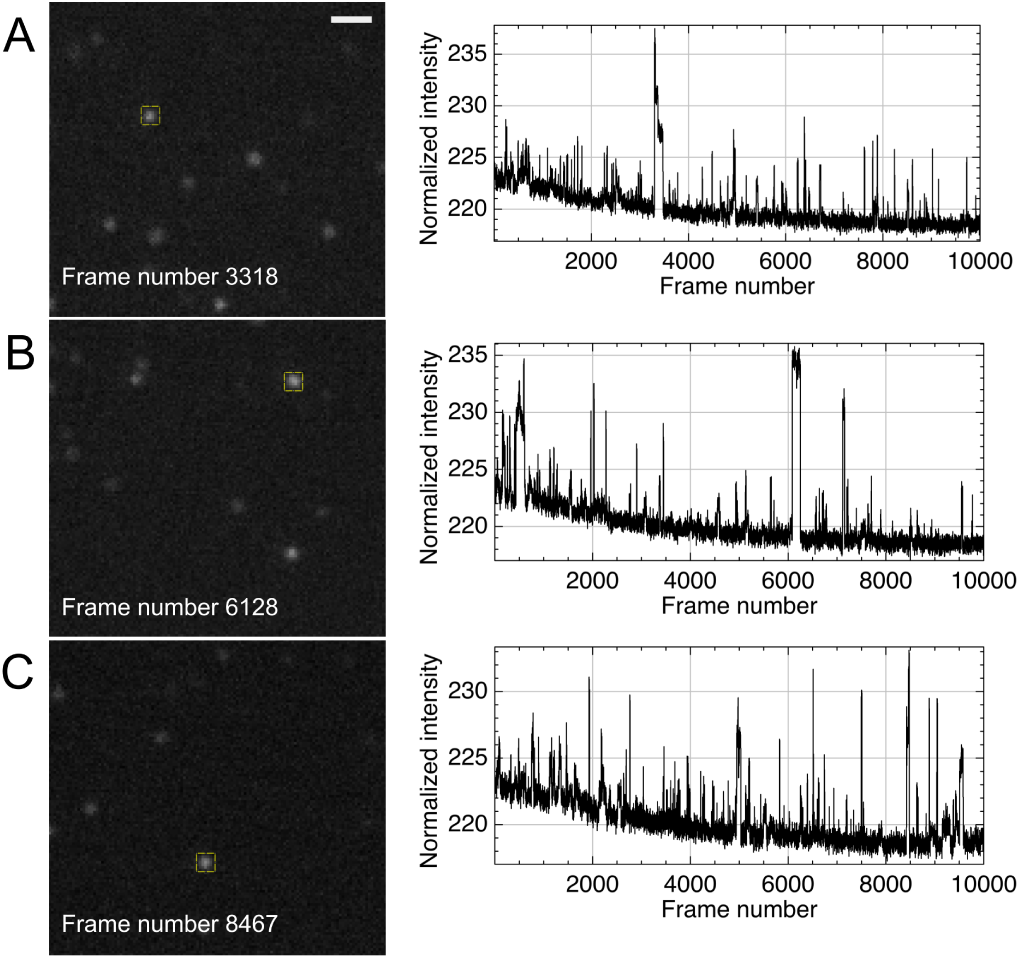
Proof of on/off switching of single molecules induced by an incoherent light source: Blinking of single molecules is shown with a Mercury arc lamp. (A-C) Three selected frames (3318, 6128 and 8467) out of 10000 frames are shown on the left side where a subset of molecules are ‘on’. On the right side, intensity profiles for three 7X7 (roughly 450X450 *nm*^2^) cross sections (yellow boxes in the images on the left) along the image stack are shown. Fluorescent bursts of single molecules can be observed as sharp peaks. Scale bar: 1 *µ*m in A and is same for B, C.

### Superresolution microscopy with OLM

Next, we wondered if single molecule positions can be used to reconstruct a superresolved image. We immunostained SYCP3 (lateral elements of SC) with Alexa Fluor 594, a structure ideal for benchmarking of super-resolution microscopes. The two strands of the synaptonemal complex are 150 nm apart and cannot be separated with a confocal microscope (Figure 3A-B). Using our setup, we generated localisation maps of SYCP3 by integrating approximately 50000 observations, each of which captured photons emitted during 150 ms of camera integration time. The two strands of the SYCP3 could be resolved with OLM (Figure 3C). A more precise contrast between confocal and OLM images is quantified using line scans (Figure 3D). On average, we detected 1000 photons per signal (Figure 3E) with a localization precision of 20 nm (Figure 3F). We achieved a Fourier Ring Correlation (FRC) resolution (Nieuwenhuizen et al., 2013) of about 90 nm (Figure 3G) in the case of SC samples.

**Figure 3.**
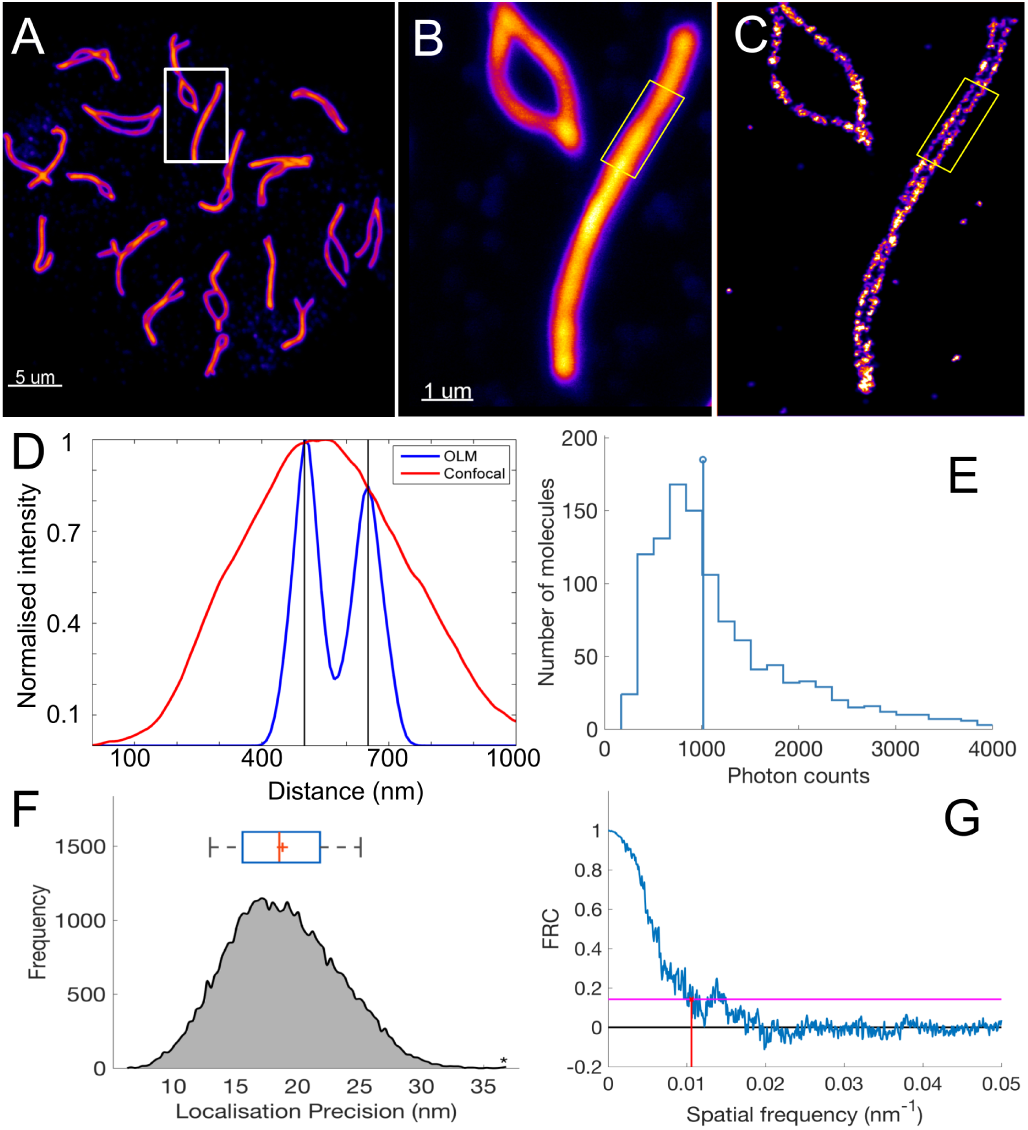
Single molecule superresolution microscopy with OLM: Pachytene chromosomes from mouse stained with an antibody against the synaptonemal complex (SC). (A) The low magnification confocal image showing the entire complement of chromosomes. (B) The boxed area in (A) shown at a higher magnification of the confocal microscope. Note that the two halves of the SC that unite the meiotic chromosome pair are not resolved. (C) The same chromosome imaged with OLM and the two halves of the SC could be resolved. (D) The contrast between OLM and confocal is further demonstrated by line scans of the boxed regions in B and C. Line scans show that OLM can resolve the two halves of the synaptonemal complex (about 150 nm separation). (E) An average number of photons (median: 1000 photons/signal) extracted from single molecules after illumination with a Mercury arc lamp. (F) An average localisation precision of 20 nm is obtained for Alexa Fluor 594. The blue box above the plot denotes the 25% and 75% quartiles while the whiskers bound 9% and 91% of the data. The red line in box denotes median while the ‘+’ sign indicates the mean. (G) FRC resolution around 90 nm is currently being achieved with OLM for SC samples. The inverse of spatial frequency (red line) provides an estimate for the resolution. The horizontal pink line indicates the 1/7 threshold of the radially combined Fourier frequencies as suggested by Nieuwenhuizen et al. (2013). Scale bar: 5 *µ*m in A. 1 *µ*m in B and is same for C.

We further compared localisation maps of SYCP3 with OLM and that of a standard high-end SMLM setup (Figure S2A-B), each of which captured photons emitted during 150 ms and 100 ms of camera integration time, respectively. On average, we detected 1000 photons per cycle, which is comparable to the photons per cycle we get with Alexa Fluor 555 with standard SMLM (Figure S2C). The setups localised individual fluorophores with an average precision of 11 nm (standard SMLM) and 18 nm (OLM) (Figure S2D). Presently, we achieve an effective FRC resolution of 67 nm for Alexa Fluor 555 when illuminated with a laser (Figure S2E), while a FRC resolution of 94 nm for Alexa Fluor 594 when illuminated with a Mercury arc lamp (Figure S2F).

### Photo re-activation of single molecules provides high signal density

We next discovered that 350-380 nm spectral peaks in the Mercury arc lamp could also be used for photoactivation of single molecules. We could make a fluorochrome blink for an extended recording of the data. A continuous exposure of the sample with the Mercury arc lamp (tested up to 12 hours) did not lead to permanent bleaching (Figure 4A-D). One of the current problems with localisation microscopy is that the high laser power tends to bleach the sample before one can record sufficient signal density to make biological inferences. In the case of OLM, it might be possible that the relatively low power and non-coherent nature of the Mercury arc lamp help in preventing the permanent bleaching of the sample.

**Figure 4.**
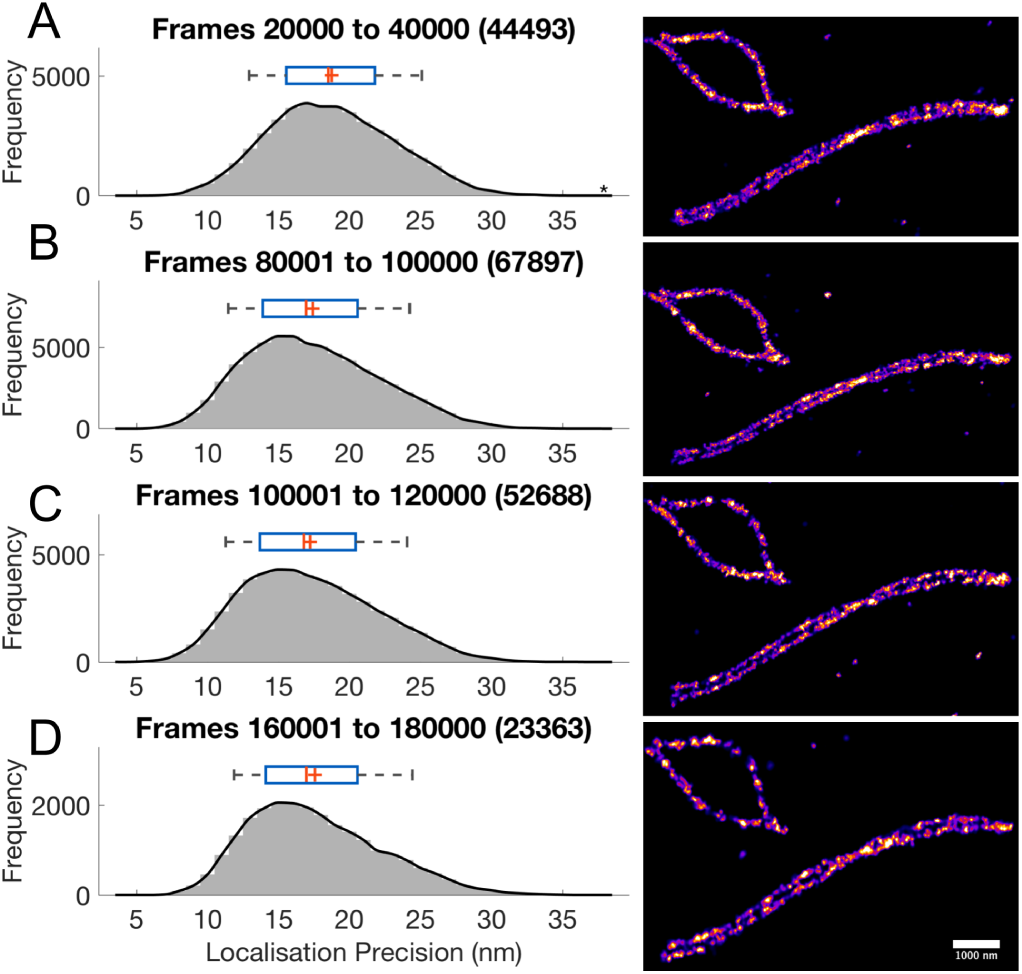
Photo re-activation of single molecules and prolonged switching of single molecules with the Mercury arc lamp. (A-D) shows independent acquisitions of SC labelled with Alexa Fluor 594. The mean localisation precision (approx. 20 nm) was consistent over the extended period of data acquisition. (A) In total, 44493 signals were acquired in an independent acquisition of 20000 frames. (B) Frames 80000-100000 with 67897 signals, (C) Next 20000 frames with 52688 signals and (D) Last 20000 frames with 23363 signals. These signals were recorded over a period of 12 hours. The blue box above each plot denotes the 25% and 75% quartiles while the whiskers bound 9% and 91% of the data. The red line in the box for each plot denotes the median while the ‘+’ is for the mean. Scale bar: 1 *µ*m in D and is same for A, B, C.

We would like to stress here that high signal density with good localisation precision and the low duty cycle are critical to reconstruct superresolution images from the single molecule data (Dempsey et al., 2011; Legant et al., 2016). For instance, we would need about 1000 signals in this diffraction limited spot to get a Nyquist resolution of under 10 nm (Ha and Tinnefeld, 2012). With indefinite blinking of Alexa Fluor 594, getting 1000 signals with good precision is easy, however, it has to be complemented with low duty cycle and higher localisation accuracy of the fluorophore.

### Photo re-activation can be used for correlative OLM and STED

Next, we observed that using DAPI excitation filter (365/30 nm), we could recover the sample after STED bleaching. Figure 5A-B shows a bleached cross section of nuclear envelope after illumination with STED acquisition. We used spectral peaks (range 350-380 nm) of the Mercury arc lamp to recover the bleached sample (Figure 5C).

**Figure 5.**
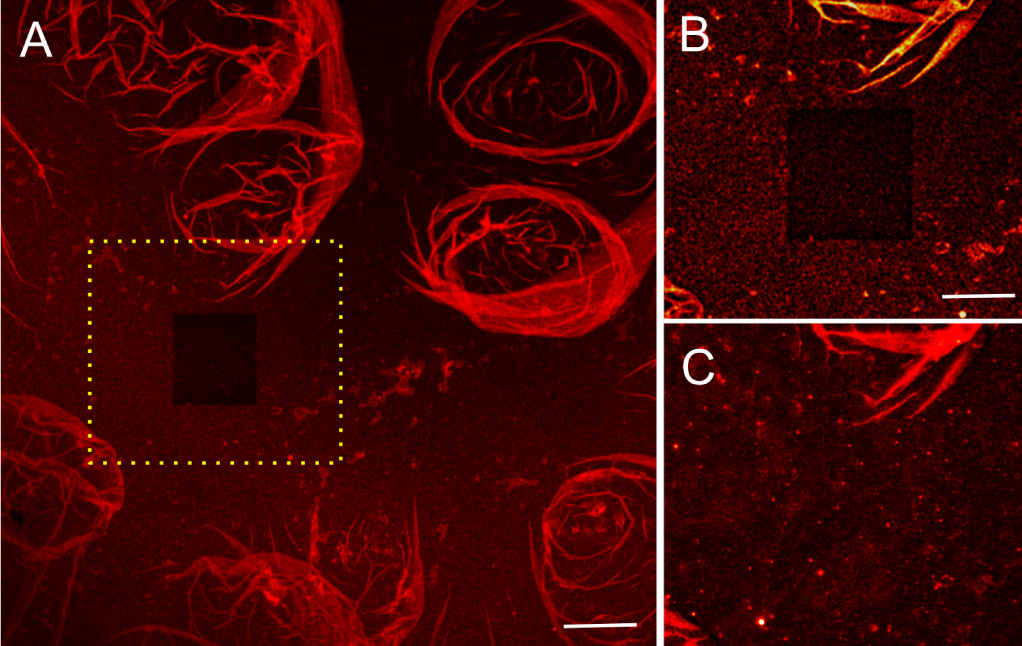
Photo re-activation of a nuclear membrane cross section using the Mercury arc lamp: (A) shows the bleached cross-section after STED imaging. (B) The section in yellow box in (A). (C) The same photobleached section as in (B) after the photoactivation. Scale bar: 2.5 *µ*m in A. 1 *µ*m in B and is same for C.

Alexa Fluor 594, due to its brightness, stable in nature, also happens to be a good STED fluorochrome. Moreover, the STED depletion laser wavelength does not excite the dye. We wondered if we can make use of the reactivation property of Alexa Fluor 594 with UV illumination to do correlative OLM and STED. We first imaged Alexa Fluor 594 in the confocal mode (Figure 6A) with Leica SP8 and then in STED mode (Figure 6D), with 660 and 775 nm depletion lasers. Next, we imaged the same region of the sample on Olympus BX 61 in the widefield mode (Figure 6B). Once bleached, we used UV illumination (365/30 nm filter) to recover the sample and then Texas Red filter to excite a subset of photo-activated molecules for OLM imaging (Figure 6E).

**Figure 6.**
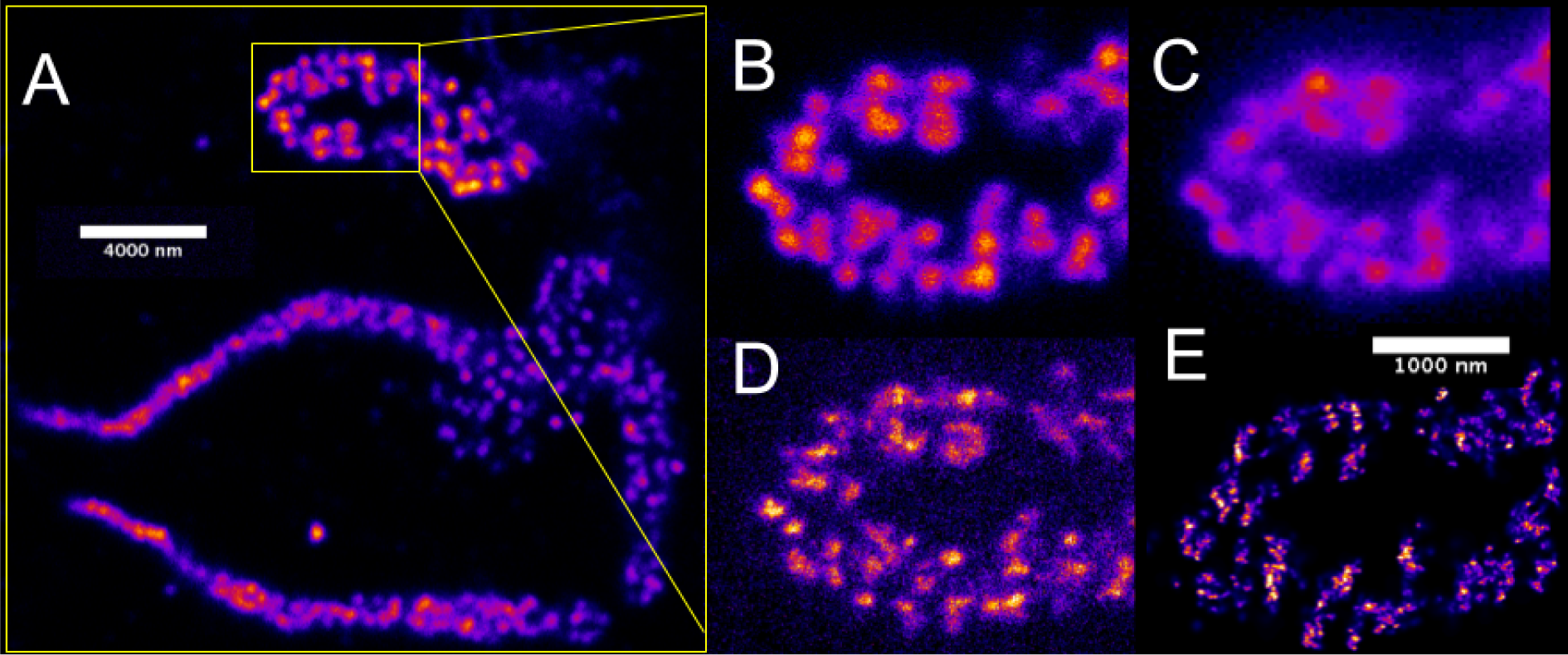
Correlative OLM and STED microscopy: Distribution of RNA-binding protein CELF1 stained with Alexa Fluor 594 on specific loops of lampbrush chromosomes (LBCs) is compared using different light microscopy techniques. (A) The low magnification confocal image is showing the entire complement of the lampbrush chromosomes. (B) The boxed area in A shown at a higher magnification using confocal (B), widefield (C), STED (D) and OLM (E). Scale bar: 4 *µ*m in A. 1 *µ*m in E and is same for B, C, D.

Lampbrush chromosomes are around 1 *µ*m thick, and due to the epi-illumination scheme of OLM, we detected many out-of-focus signals. We attribute the minor structural differences in OLM and STED images to this factor. The optical sectioning drawback of OLM can be easily overcome by using a TIRF objective. In this regard, depending upon the thickness of the sample the general optimisation rules for OLM might change.

## Discussion

In this proof-of-principle study, we have shown that highdensity superresolution images can be obtained using a Mercury arc lamp and a conventional epifluorescence microscope setup. The configuration presented here can be easily extended to other light sources such LEDs or metal Halide lamps, given that they can provide a minimum threshold energy within the excitation bandwidth of the fluorophore (Lichtman and Conchello, 2005). To note, the metal Halide lamps provide a better alternative to the Mercury arc lamps as they have a brighter intensity between 460-520 nm range, a more uniform field of illumination, and allow more control of the lamp power (Figure S3). Moreover, they produce less heat than the Xenon and Mercury arc lamps, and no bulb alignment is required.

Once the illumination is optimised, care must be taken that different fluorescent moieties have a low duty cycle in addition to having high photon yield and photostability in order to achieve high-resolution single molecule reconstructions (Dempsey et al., 2011). Limited signal density due to photobleaching of fluorophores has been a central problem with most single-molecule superresolution microscopy techniques, so far. The prolonged reactivation of Alexa Fluor 594 is particularly helpful to re-use the same sample for multiple measurements and perform imaging on different setups. The prolonged blinking of a fluorophore can be further utilised for single particle tracking experiments (Manley et al., 2008; Gahlmann and Moerner, 2014; Balzarotti et al., 2017). The reactivation with UV illumination is similar to that of Alexa Fluor 647 in a redox buffer but in the present case using light instead of chemicals. It might be possible that lasers bleach the fluorochrome permanently and faster, which might not be the reason in the case of an incoherent light source. A proper calibration for different fluorophores with both lamp and laser is suggested.

We were also able to perform STED and OLM on the same biological sample with comparable resolution, confirming the high-resolution setting of the setup. Prolongated blinking with deep blue illumination was key to achieve a high signal density and subsequently the high resolution. This comes from the fact that the final resolution of an image depends not only on the localisation precision but also on the total number of independently localised signals (Legant et al., 2016).

At this point, we wish to re-emphasize that the blinking phenomenon can be observed with ProLong Diamond. This simplifies the need to prepare special buffers consisting of the classical redox cocktail required for dSTORM and other related superresolution techniques (Heilemann et al., 2008). As the composition of Prolong Diamond is not known, it was difficult to narrow down the minimal requirements for blinking to happen at this stage of the study. This results hints at a conformational explanation of blinking events (Baddeley et al., 2009; Estévez-Torres et al., 2009), in addition to cycles of *H*^+^ binding and release, as stated before (Fölling et al., 2008; Żurek-Biesiada et al., 2015, 2016).

The two main challenges associated with OLM and SMLM techniques, in general, are the stage/sample drift and optical sectioning (Betzig et al., 2006; Juette et al., 2008). Due to sample drift (correction only up to 50 nm) and high duty cycle of Alexa Fluor 594, we could not resolve the individual components of the NPCs, which are only 20-40 nm apart. In the case of OLM, the majority of drift stemmed from heat generated by the Mercury arc lamp, stage movement and objective holder. Stabilising the whole microscope for at least one hour before the measurement significantly helps in minimizing the drift. For good optical sectioning, we made use of very thin samples. For thick samples, TIRF objectives can be used to achieve a better signal-to-noise ratio. Furthermore, camera readout noise (CCD vs EMCCD) can be a significant factor for the fluorophores with a poor photon count.

The overall cost of the setup is low and comes with minimal implementation effort. Most of other high-resolution imaging techniques require the use of relatively high power lasers, expensive objective lens (high-NA), very accurate piezoelectric stages and high-end cameras. Lasers are difficult to align, and misalignment often produces artefacts that can be difficult to recognise if the prior information about the sample is not available (Prakash, 2017b). Moreover, the technical aspect of the implementation procedures makes these setups a difficult access to many biologists. We hope that OLM paves the way for a simple and low-cost high-resolution microscopy implementation especially in emerging countries with limited science budget.

We think that the accessibility of the method and its relative easy use can democratise superresolution imaging and make it an everyday use in molecular biology studies. Last but not least, various photophysical observation such as indefinite blinking due to photoreactivation and non-permanent bleaching of fluorophores with deep blue illumination prompts for more thorough investigation on the photophysics of a fluorophore (Lippincott-Schwartz et al., 2003; Lippincott-Schwartz and Patterson, 2008) and various mechanisms that can make a fluorophore blink (Vogelsang et al., 2010).

### Materials and Methods

All experimental procedures were performed in compliance with ethical regulations and approved by the IACUC of the Carnegie Institution for Science. The meiotic spreads in mouse oocyte, nuclear pore complex samples, and the lampbrush chromosome samples were kindly provided by Joseph G Gall, Zehra Nizami, and Safia Maliki. The details about the sample preparation can be found in the following articles (Susiarjo et al., 2009; Malki and Bortvin, 2017; Prakash, 2017c; Gall and Nizami, 2016; Shi et al., 2017).

#### Immunofluorescence

Sample coverslips were washed for 5 min in (PBS, 0.05% Triton X-100) and incubated for 1 hour at room temperature with blocking solution (PBS, 0.05% Triton X-100, 10% Normal serum). We applied rabbit polyclonal anti-SYCP3 (1:1000, ab15092) in the antibody solution (PBS, 0.05% Triton X-100, 10% Normal serum) and incubated for 4 hours at room temperature. We washed coverslips in (PBS, 0.1% Triton X-100) for 5 min, followed by two 5 min washes in (PBS, 0.05% Triton X-100). We detected primary antibodies by incubating coverslips for 1 hour 30 at room temperature with Alexa Fluor 594 donkey anti-rabbit (Invitrogen, R37119). After incubation, coverslips were washed in PBS (0.1% Triton X-100) for 5 min, followed by two 5 min washes in PBS (0.05% Triton X-100). This protocol was provided by Safia Malki.

#### Imaging media

We rinsed coverslips in water and used ProLong Diamond (ThermoFIsher, P36970) as the anti-fading mounting solution. The refractive index of the oil used was 1.518. We wish to emphasise here that ProLong Diamond works successfully for both STED and OLM/SMLM microscopy with Alexa Fluor 594. For two or more colours imaging, a more complex and optimised imaging buffer might be required.

#### Microscopy

Confocal and STED images were obtained using a Leica TCS SP8 microscope with 592-nm, 660-nm, and 775-nm depletion lasers. OLM images were acquired with an Olympus BX61 microscope equipped with 100x / NA 1.4 Oil objective (Olympus UPlanSApo) and a Hamamatsu CCD camera (C4742-95). The effective pixel size was 64.5 nm in the sample plane. Illumination was done using a Mercury arc lamp (100W, Ushio USH-103D), as is commonly used in the conventional epifluorescence microscopes. The filter cube consisted of the following excitation filters: Texas Red (560/40 nm) and DAPI (365/30), emission filter (590/40 nm) and a dichromatic mirror. All the filters were bought from Semrock. The microscope was placed on a simple table with no stabilisation or control for vibration (Figure S4), so there was a considerable drift of the sample during the measurement. Due to reactivation of Alexa 594 with DAPI excitation filter, we could record a high number of frames, which provided us with the luxury to discard the frames with significant drift. The frames with significantly less drift were corrected using post-acquisition drift correction algorithms as described in Prakash (2017b, 2016).

#### Data acquisition

A few parameters such as bleaching time, the number of signals per frame, camera integration time and the total number of final frames need to be optimised before acquiring the data. In our case, 10-20 minutes of pre-bleaching with Texas Red excitation filter is required before molecules started to blink. The pre-bleaching step further helps to minimise autofluorescence which was minimal in the case of flat samples like synaptonemal complex (SC). Once blinking started, we adjusted the lamp power to have 10-20 signals per frame in a cross section of 25 *µm*^2^. Only a few signals per frame help to optically isolate the molecules from each other and obtain high-resolution images. We found approximately 20000 frames with 50000 localizations to be enough to reconstruct high-density images of SC. An integration time of 100-150 ms was sufficient for a good signal-to-noise ratio. Finally, to speed up the acquisition time and save data storage space, the imaging area can be restricted to a region of interest.

#### Data reconstruction and visualisation

After data acquisition, the coordinates of the single signals need to be precisely determined. Figure S1 presents an overview of various steps required to reconstruct a highly resolved image from a stack comprising a large number of images, where each image contains only a few signals.

As the first step, single molecule signals in each frame need to be separated from the background. The background varies throughout the image stacks, but the variation in the background between the consecutive images is relatively negligible. We used this strategy to estimate the background for a given frame based on the information from the previous images. The noise in the image can further make it difficult to determine the coordinates of the signal precisely. The error in the measured intensity *N* is the square root of the measured value 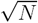, following a Poisson distribution. This error can become significant when the background signal is high and non-uniform. In the present case, we estimated a background map for each image by averaging previous ten frames and then subtracting from the image to get the difference image. Next, a median filter was applied to the difference image to get the signals above an empirically determined threshold.

In the second step, we extracted the local maxima in each frame and select the corresponding regions-of-interest (ROIs) of the signal (Grull et al., 2011; Prakash, 2017b). The centre of each signal is precisely determined by a statistical fit approximately the ideal PSF (Airy-function) with a Gaussian. In cases, where the sample background and camera noise are minimal, the fitted position can be estimated with a precision of 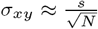, where *s* is the standard deviation of the Gaussian and *N* is the total number of photons detected (Betzig et al., 2006; Thompson et al., 2002). An updated formula by Mortensen et al. (2010) and Stallinga and Rieger (2012) provides a good approximation to calculate the localisation precision for samples with significant background and readout noise.

In our setup, some signals stay ‘on’ for a time longer than the camera integration time. If the signal appears in the consecutive frames, then it can be removed or merged to prevent recounting of the same signal. In the present case, we combined the signals in successive frames if the signals were within the localisation precision of the first signal. For visualisation, each signal position was blurred with a Gaussian distribution of standard deviation equal to the mean distance to the next 20 nearest neighbour molecule positions. The algorithm has been described previously in (Prakash et al., 2015; Kaufmann et al., 2012).

Publicly available SMLM data reconstruction softwares such as ThunderSTORM (Ovesný et al., 2014), QuickPALM (Henriques et al., 2010) or rapidSTORM (Wolter et al., 2012) can be used to carry out the above mentioned analysis.

## Acknowledgements

K.P. acknowledges Joseph G. Gall for the financial support. K.P. further acknowledges Joseph G. Gall, Zehra Nizami, Safia Malki, Yixian Zheng and Mahmud Siddiqi for the biological samples, help with microscopy measurements and useful discussions. K.P. thanks David Fournier, Wioleta Dudka and Miguel Andrade-Navarro for proofreading the manuscript.

## Conflict of interest

The author declare no conflict of interest.

## Supplemental Material

**High-density super-resolution microscopy with an incoherent light source and a conventional epifluorescence microscope setup**

**Figure S1.**
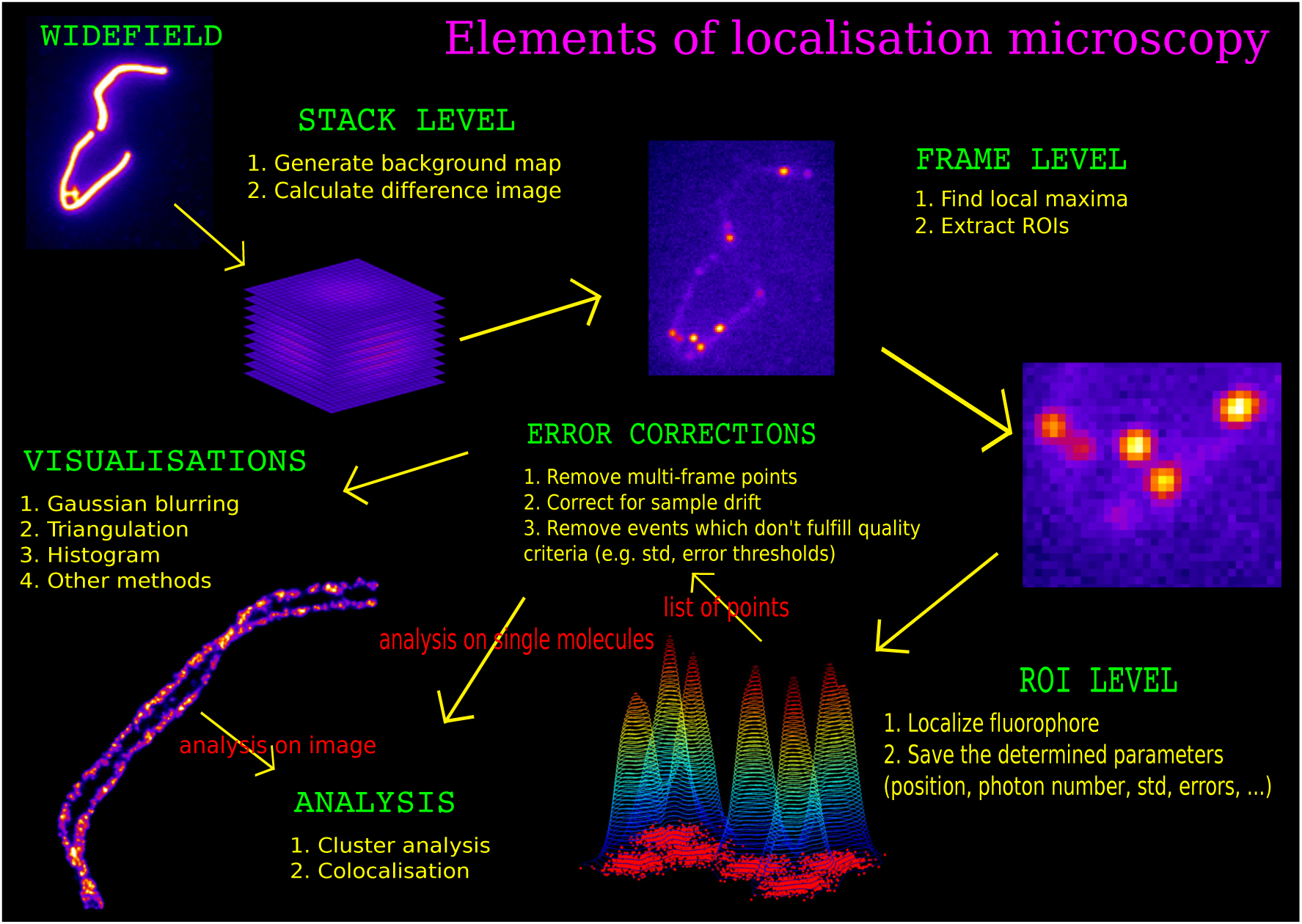
OLM data reconstruction and analysis flowchart (Prakash, 2017a). See Methods and Material section for details.

**Figure S2.**
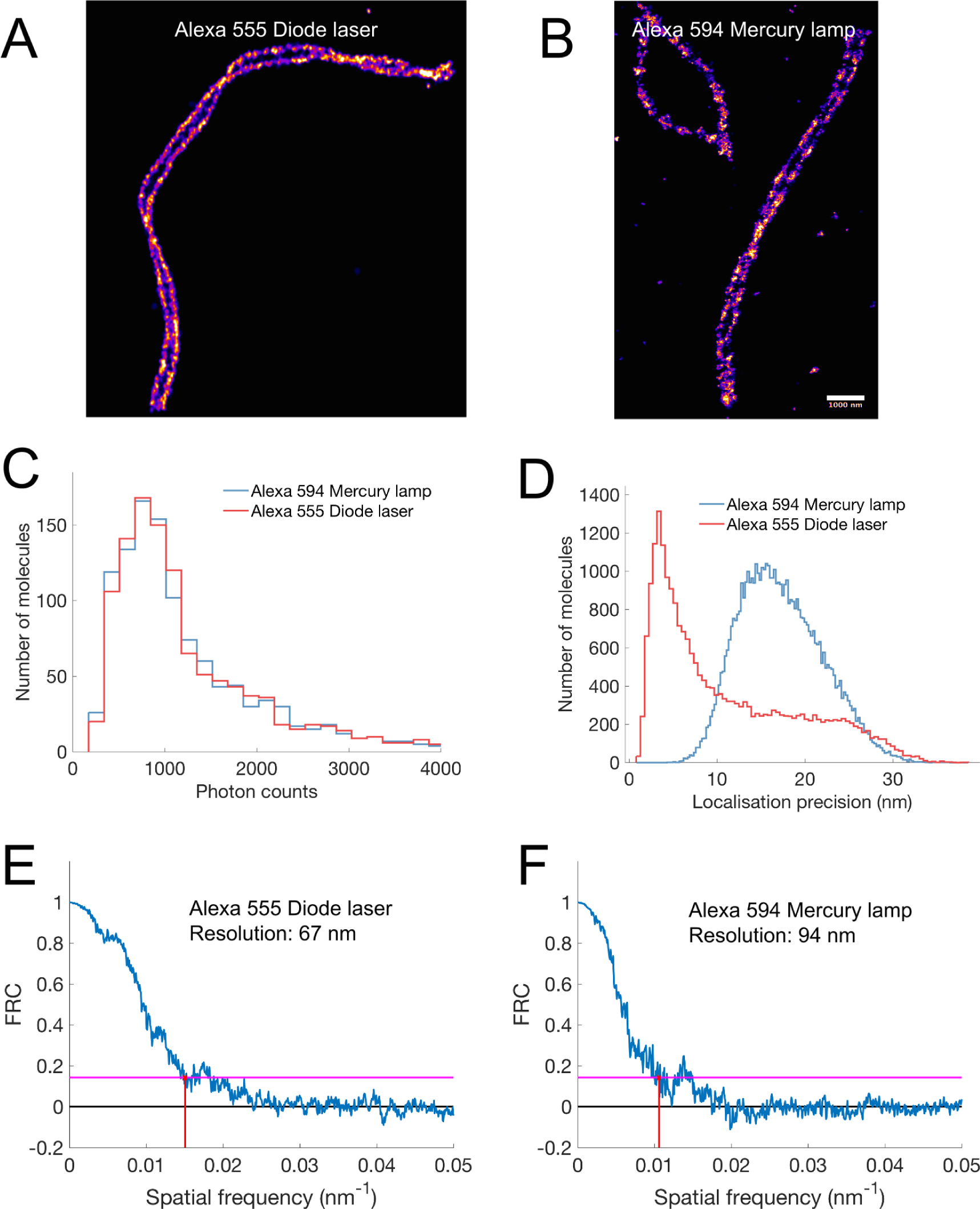
Comparision of laser-based SMLM and lamp-based OLM: (A) SMLM image of SYCP3 labelled with Alexa Fluor 555 (Prakash et al., 2015). Image acquired with a high-end wide-field setup consisting of a diode laser, a very sensitive camera, piezo stage and a high NA objective. (B) OLM image of SYCP3 labelled with Alexa Fluor 555. The image is acquired with Olympus BX61. The setup consists of an incoherent light source, a simple CCD camera with an ordinary stage. (C) Comparison of photons emitted by single molecules using these two setups. (D) Comparison of the two configurations using localisation precision. (E-F) FRC resolution for Alexa Fluor 555 (SMLM) and Alexa Fluor 594 (OLM).

**Figure S3.**
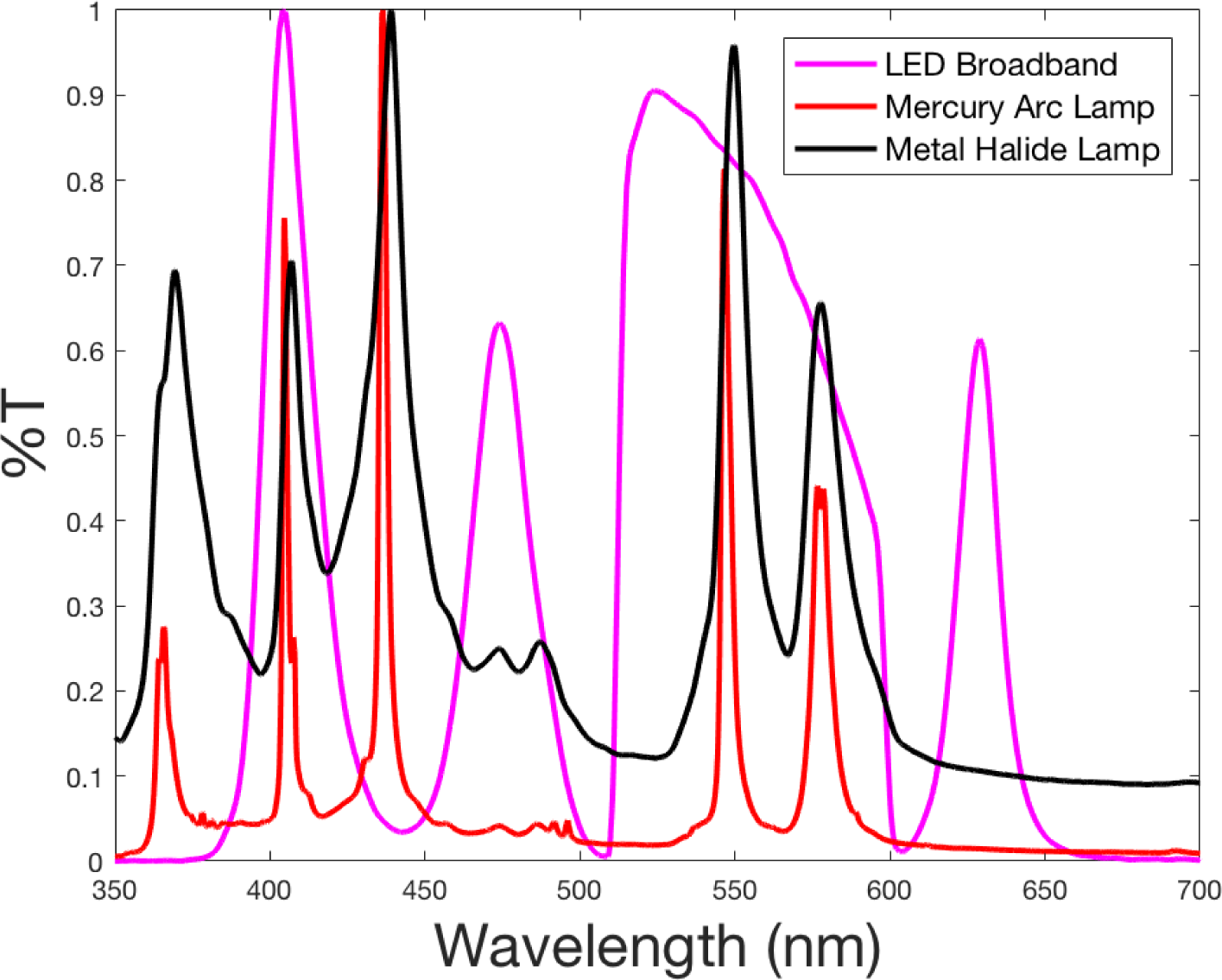
A comparison of spectra of a standard LED broadband, Mercury arc lamp and metal Halide lamp. Both LEDs and metal Halide lamps provide more uniform illumination than the Mercury arc lamps and can be good alternatives.

**Figure S4.**
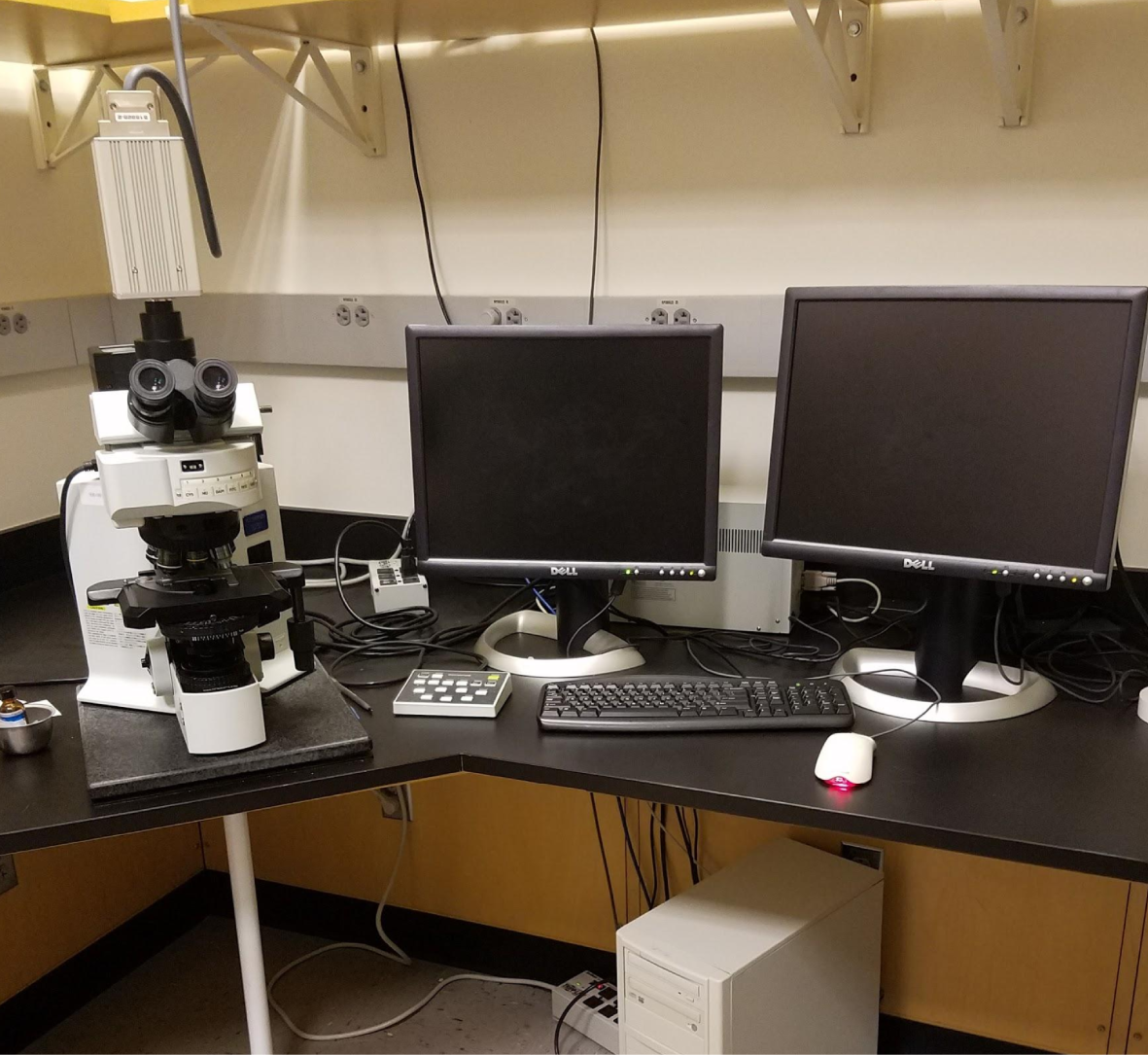
Olympus BX 61 as used for OLM measurements.

## References

Abbe, E. (1873). Beiträge zur Theorie des Mikroskops und der mikroskopischen Wahrmehmung. Archiv für Mikroskopische Anatomie, 9:413–420.

Baddeley, D., Jayasinghe, I. D., Cremer, C., Cannell, M. B., and Soeller, C. (2009). Light-induced dark states of organic fluochromes enable 30 nm resolution imaging in standard media. Biophysical journal, 96(2):L22–L24.

Balzarotti, F., Eilers, Y., Gwosch, K. C., Gynnå, A. H., Westphal, V., Stefani, F. D., Elf, J., and Hell, S. W. (2017). Nanometer resolution imaging and tracking of fluorescent molecules with minimal photon fluxes. Science, 355(6325):606–612.

Betzig, E., Patterson, G. H., Sougrat, R., Lindwasser, O. W., Olenych, S., Bonifacino, J. S., Davidson, M. W., Lippincott-Schwartz, J., and Hess, H. F. (2006). Imaging intracellular fluorescent proteins at nanometer resolution. Science, 313(5793):1642–1645.

Betzig, E. and Trautman, J. K. (1992). Near-field optics: microscopy, spectroscopy, and surface modification beyond the diffraction limit. Science, 257(5067):189–196.

Born, M. and Wolf, E. (1980). *Principles of optics: electromagnetic theory of propagation, interference and diffraction of light*. Elsevier.

Chiu, H. and Quake, S. (1999). Single-molecule fluorescence observed with mercury lamp illumination. Biotechniques, 27(5):1008–1014.

Cremer, C. and Masters, B. R. (2013). Resolution enhancement techniques in microscopy. The European Physical Journal H, 38(3):281–344.

Dempsey, G. T., Vaughan, J. C., Chen, K. H., Bates, M., and Zhuang, X. (2011). Evaluation of fluorophores for optimal performance in localization-based super-resolution imaging. Nature methods, 8(12):1027–1036.

Dertinger, T., Colyer, R., Iyer, G., Weiss, S., and Enderlein, J. (2009). Fast, background-free, 3d super-resolution optical fluctuation imaging (sofi). Proceedings of the National Academy of Sciences, 106(52):22287–22292.

Estévez-Torres, A., Crozatier, C., Diguet, A., Hara, T., Saito, H., Yoshikawa, K., and Baigl, D. (2009). Sequence-independent and reversible photocontrol of transcription/expression systems using a photosensitive nucleic acid binder. Proceedings of the National Academy of Sciences, 106(30):12219–12223.

Fölling, J., Bossi, M., Bock, H., Medda, R., Wurm, C. A., Hein, B., Jakobs, S., Eggeling, C., and Hell, S. W. (2008). Fluorescence nanoscopy by ground-state depletion and single-molecule return. Nature methods, 5(11):943–945.

Gahlmann, A. and Moerner, W. (2014). Exploring bacterial cell biology with single-molecule tracking and super-resolution imaging. Nature Reviews Microbiology, 12(1):9–22.

Gall, J. G. and Nizami, Z. F. (2016). Isolation of giant lampbrush chromosomes from living oocytes of frogs and salamanders. JoVE (Journal of Visualized Experiments), (118):e54103–e54103.

Gerhardt, I., Mai, L., Lamas-Linares, A., and Kurtsiefer, C. (2011). Detection of single molecules illuminated by a light-emitting diode. Sensors, 11(1):905–916.

Grull, F., Kirchgessner, M., Kaufmann, R., Hausmann, M., and Kebschull, U. (2011). Accelerating image analysis for localization microscopy with fpgas. In Field Programmable Logic and Applications (FPL), 2011 International Conference on, pages 1–5. IEEE.

Gustafsson, M. G. (2000). Surpassing the lateral resolution limit by a factor of two using structured illumination microscopy. Journal of microscopy, 198(2):82–87.

Gustafsson, N., Culley, S., Ashdown, G., Owen, D. M., Pereira, P. M., and Henriques, R. (2016). Fast live-cell conventional fluorophore nanoscopy with imagej through super-resolution radial fluctuations. Nature Communications, 7.

Ha, T. and Tinnefeld, P. (2012). Photophysics of fluorescent probes for single-molecule biophysics and super-resolution imaging. Annual review of physical chemistry, 63:595–617.

Heilemann, M., Van De Linde, S., Schüttpelz, M., Kasper, R., Seefeldt, B., Mukherjee, A., Tinnefeld, P., and Sauer, M. (2008). Subdiffraction-resolution fluorescence imaging with conventional fluorescent probes. Angewandte Chemie International Edition, 47(33):6172–6176.

Heintzmann, R. and Cremer, C. (1999). Laterally modulated excitation microscopy: improvement of resolution by using a diffraction grating. In BiOS Europe’98, pages 185–196. International Society for Optics and Photonics.

Hell, S. W. and Wichmann, J. (1994). Breaking the diffraction resolution limit by stimulated emission: stimulated-emission-depletion fluorescence microscopy. Optics letters, 19(11):780–782.

Henriques, R., Lelek, M., Fornasiero, E. F., Valtorta, F., Zimmer, C., and Mhlanga, M. M. (2010). Quickpalm: 3d real-time photoactivation nanoscopy image processing in imagej. Nature methods, 7(5):339–340.

Hess, S. T., Girirajan, T. P., and Mason, M. D. (2006). Ultra-high resolution imaging by fluorescence photoactivation localization microscopy. Biophysical journal, 91(11):4258.

Juette, M. F., Gould, T. J., Lessard, M. D., Mlodzianoski, M. J., Nagpure, B. S., Bennett, B. T., Hess, S. T., and Bewersdorf, J. (2008). Three-dimensional sub–100 nm resolution fluorescence microscopy of thick samples. Nature methods, 5(6):527–529.

Kaufmann, R., Piontek, J., Grüll, F., Kirchgessner, M., Rossa, J., Wolburg, H., Blasig, I. E., and Cremer, C. (2012). Visualization and quantitative analysis of reconstituted tight junctions using localization microscopy. PloS one, 7(2):e31128.

Keller, P. J., Schmidt, A. D., Santella, A., Khairy, K., Bao, Z., Wittbrodt, J., and Stelzer, E. H. (2010). Fast, high-contrast imaging of animal development with scanned light sheet-based structured-illumination microscopy. Nature methods, 7(8):637–642.

Legant, W. R., Shao, L., Grimm, J. B., Brown, T. A., Milkie, D. E., Avants, B. B., Lavis, L. D., and Betzig, E. (2016). High-density three-dimensional localization microscopy across large volumes. Nature methods, 13(4):359–365.

Lichtman, J. W. and Conchello, J.-A. (2005). Fluorescence microscopy. Nature methods, 2(12):910–919.

Lidke, K., Rieger, B., Jovin, T., and Heintzmann, R. (2005). Superresolution by localization of quantum dots using blinking statistics. Optics express, 13(18):7052–7062.

Lippincott-Schwartz, J., Altan-Bonnet, N., and Patterson, G. H. (2003). Photobleaching and photoactivation: following protein dynamics in living cells. Nature Cell Biology, page S7.

Lippincott-Schwartz, J. and Patterson, G. H. (2008). Fluorescent proteins for photoactivation experiments. Methods in cell biology, 85:45–61.

Löschberger, A., van de Linde, S., Dabauvalle, M.-C., Rieger, B., Heilemann, M., Krohne, G., and Sauer, M. (2012). Superresolution imaging visualizes the eightfold symmetry of gp210 proteins around the nuclear pore complex and resolves the central channel with nanometer resolution. J Cell Sci, 125(3):570–575.

Malki, S. and Bortvin, A. (2017). Epigenetics and transposon control in the mammalian germline. In Epigenetics in Human Reproduction and Development, pages 1–33.

Manley, S., Gillette, J. M., Patterson, G. H., Shroff, H., Hess, H. F., Betzig, E., and Lippincott-Schwartz, J. (2008). High-density mapping of single-molecule trajectories with photoactivated localization microscopy. Nature methods, 5(2):155–157.

Mortensen, K. I., Churchman, L. S., Spudich, J. A., and Flyvbjerg, H. (2010). Optimized localization analysis for single-molecule tracking and super-resolution microscopy. Nature methods, 7(5):377–381.

Nieuwenhuizen, R. P., Lidke, K. A., Bates, M., Puig, D. L., Grünwald, D., Stallinga, S., and Rieger, B. (2013). Measuring image resolution in optical nanoscopy. Nature methods, 10(6):557–562.

Ovesný, M., Křížek, P., Borkovec, J., Švindrych, Z., and Hagen, G. M. (2014). Thunderstorm: a comprehensive imagej plug-in for palm and storm data analysis and super-resolution imaging. Bioinformatics, 30(16):2389–2390.

Peyser, L. A., Vinson, A. E., Bartko, A. P., and Dickson, R. M. (2001). Photoactivated fluorescence from individual silver nanoclusters. Science, 291(5501):103–106.

Prakash, K. (2016). *The periodic and dynamic structure of chromatin*. PhD thesis, Heidelberg University.

Prakash, K. (2017a). *Chromatin Architecture: Advances From High-resolution Single Molecule DNA Imaging*. Springer.

Prakash, K. (2017b). Investigating chromatin organisation using single molecule localisation microscopy. In Chromatin Architecture, pages 25–61. Springer.

Prakash, K. (2017c). Periodic and symmetric organisation of meiotic chromosomes. In Chromatin Architecture, pages 105–133. Springer International Publishing.

Prakash, K., Fournier, D., Redl, S., Best, G., Borsos, M., Tiwari V. K., Tachibana-Konwalski, K., Ketting, R. F., Parekh, S. H., Cremer, C., et al. (2015). Superresolution imaging reveals structurally distinct periodic patterns of chromatin along pachytene chromosomes. Proceedings of the National Academy of Sciences, 112(47):14635–14640.

Rayleigh, L. (1896). Xv. on the theory of optical images, with special reference to the microscope. The London, Edinburgh, and Dublin Philosophical Magazine and Journal of Science, 42(255):167–195.

Rust, M. J., Bates, M., and Zhuang, X. (2006). Stochastic optical reconstruction microscopy (storm) provides sub-diffraction-limit image resolution. Nature methods, 3(10):793.

Schoen, I., Ries, J., Klotzsch, E., Ewers, H., and Vogel, V. (2011). Binding-activated localization microscopy of dna structures. Nano letters, 11(9):4008–4011.

Sheppard, C. and Wilson, T. (1981). Effects of high angles of convergence on v (z) in the scanning acoustic microscope. Applied Physics Letters, 38(11):858–859.

Shi, K. Y., Mori, E., Nizami, Z. F., Lin, Y., Kato, M., Xiang, S., Wu, L. C., Ding, M., Yu, Y., Gall, J. G., et al. (2017). Toxic prn poly-dipeptides encoded by the c9orf72 repeat expansion block nuclear import and export. Proceedings of the National Academy of Sciences, page 201620293.

Stallinga, S. and Rieger, B. (2012). The effect of background on localization uncertainty in single emitter imaging. In Biomedical Imaging (ISBI), 2012 9th IEEE International Symposium on, pages 988–991. IEEE.

Susiarjo, M., Rubio, C., and Hunt, P. (2009). Analyzing mammalian female meiosis. In *Meiosis*, pages 339–354. Springer.

Szczurek, A. T., Prakash, K., Lee, H.-K., Żurek-Biesiada, D. J., Best, G., Hagmann, M., Dobrucki, J. W., Cremer, C., and Birk, U. (2014). Single molecule localization microscopy of the distribution of chromatin using hoechst and dapi fluorescent probes. Nucleus, 5(4):331–340.

Thompson, R. E., Larson, D. R., and Webb, W. W. (2002). Precise nanometer localization analysis for individual fluorescent probes. Biophysical journal, 82(5):2775–2783.

van’t Hoff, M., de Sars, V., and Oheim, M. (2008). A programmable light engine for quantitative single molecule tirf and hilo imaging. Optics express, 16(22):18495–18504.

Vogelsang, J., Steinhauer, C., Forthmann, C., Stein, I. H., Person-Skegro, B., Cordes, T., and Tinnefeld, P. (2010). Make them blink: Probes for super-resolution microscopy. ChemPhysChem, 11(12):2475–2490.

Willig, K. I., Rizzoli, S. O., Westphal, V., Jahn, R., and Hell, S. W. (2006). Sted microscopy reveals that synaptotagmin remains clustered after synaptic vesicle exocytosis. Nature, 440(7086):935–939.

Wolter, S., Löschberger, A., Holm, T., Aufmkolk, S., Dabauvalle, M.-C., Van De Linde, S., and Sauer, M. (2012). rapidstorm: accurate, fast open-source software for localization microscopy. Nature methods, 9(11):1040–1041.

Zipfel, W. R., Williams, R. M., and Webb, W. W. (2003). Nonlinear magic: multiphoton microscopy in the biosciences. Nature biotechnology, 21(11):1369–1377.

Żurek-Biesiada, D., Szczurek, A. T., Prakash, K., Best, G., Mohana, G. K., Lee, H.-K., Roignant, J.-Y., Dobrucki, J. W., Cremer, C., and Birk, U. (2016). Quantitative super-resolution localization microscopy of dna in situ using vybrant(r) dyecycle violet fluorescent probe. Data in Brief, 7:157–171.

Żurek-Biesiada, D., Szczurek, A. T., Prakash, K., Mohana, G. K., Lee, H.-K., Roignant, J.-Y., Birk, U., Dobrucki, J. W., and Cremer, C. (2015). Localization microscopy of dna in situ using vybrant(r) dyecycle violet fluorescent probe: A new approach to study nuclear nanostructure at single molecule resolution. Experimental cell research.

